# Locked Nucleic Acid Stabilized Liquid Crystalline Phases

**DOI:** 10.1101/2025.10.16.682820

**Authors:** Sineth G. Kodikara, Samuel Sprunt, Hamza Balci

## Abstract

Liquid crystalline (LC) phases formed by gapped DNA (GDNA) constructs, where two rigid duplexes are connected with a flexible single stranded linker, offer a versatile platform to investigate interactions between DNA molecules. Base pairs containing a locked nucleic acid (LNA-DNA or LNA-LNA pairs) are generally more stable compared to DNA-DNA pairs due to enhanced hydrogen bonding and/or attractive base stacking interactions. In concentrated solutions of GDNA constructs, the stability of terminal base pairs and the base stacking interactions between neighboring duplexes are critical for the formation of a bilayer smectic phase. By using temperature-resolved synchrotron small-angle X-ray scattering (SAXS) measurements, we quantified the impact of single LNA modification of terminal base pairs on the thermal stability of smectic LC phases. We observe that LNA-DNA terminal AT base pairing (A+T) increases the stability of the bilayer smectic phase by ∼9-18 °C relative to DNA-DNA pairing at the same duplex concentrations. While relatively large, this increase is still significantly less than the up to ∼30 °C increase observed when AT DNA-DNA pairs are replaced by GC pairs, suggesting the stacking interactions between A+T LNA-DNA base pairs are significantly weaker than those between unmodified GC base pairs. Our study illustrates the sensitivity of LC ordering in dense DNA solutions to a single nucleotide modification and demonstrates that LNA modifications can provide a new mechanism for tuning the stability of nucleic acid-based materials.

## Introduction

Liquid crystalline (LC) materials formed by concentrated solutions of gapped DNA (GDNA) have proven to be a versatile system to investigate the interactions between DNA molecules ^1–3^. By connecting two short and rigid segments (double stranded DNA--dsDNA) with a flexible linker (single stranded DNA--ssDNA), GDNA molecules create a system where these interactions and entropic considerations, due to segregation of the flexible linkers to a specific region, drive formation of layered (smectic) LC phases (Fig. 1). The confinement of the dsDNA molecules to layers further facilitates side-by-side and end-to-end interactions between neighboring duplexes. To illustrate, modifying the terminal base pairs (bp) of the duplexes of a GDNA construct from AT to GC was demonstrated to increase the thermal stability of the resulting bilayer smectic phase by ∼30 °C, which was primarily attributed to more efficient end-to-end stacking between GC - GC compared to AT - AT base pairs ^4^. In another example, the stability of in-plane positional order was found to be strongly dependent on divalent cation concentration, with melting temperature increasing for concentrations up to ∼30 mM MgCl_2_ and decreasing beyond that ^5^. This was attributed to effective charge shielding and attractive interactions due to Mg^2+^ ions and their distribution along the major and minor grooves. In both examples it was possible to determine the thermal melting temperatures and stabilities of associated phases by monitoring the temperature evolution of small angle X-ray scattering (SAXS) profiles.

**Figure 1.**
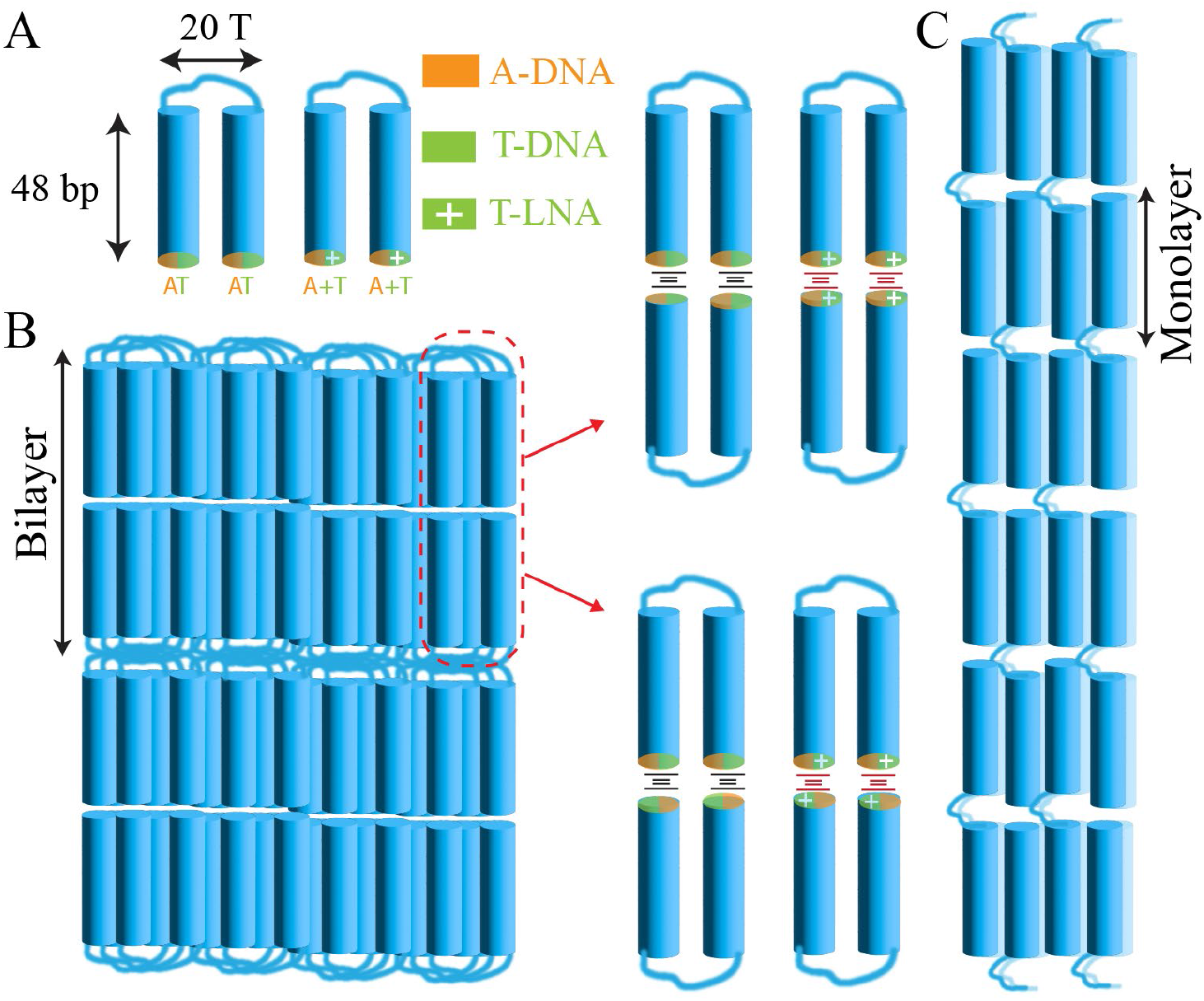
Schematics of 48-20T-48 GDNA constructs with or without LNA modifications at the terminal base pairs (A+T - A+T vs AT - AT). These constructs have 48 bp long duplex arms connected with a flexible 20T long linker (“gap”). The constructs are identical except the terminal base pairs of the duplex arms. (B) Schematics showing the bilayer order for the folded conformation of the GDNA constructs and some of the potential stacking interactions between the terminal base pairs. A schematic of the bilayer for the unfolded conformation is shown in Supporting Fig. S1. (C) Schematic of the monolayer order.

Locked nucleic acid (LNA) is a modified nucleic acid where the ribose sugar ring is “locked” in a specific conformation (C3’-endo) via a methyl bridge between the 2′-oxygen and 4′-carbon ^6,7^. This reduced flexibility results in tighter and more specific hybridization with complementary DNA or RNA sequences, resulting in 3-8 °C increase in thermal melting temperature per LNA modification ^8,9^. The enhanced stability has been attributed primarily to stronger hydrogen bonds or elevated stacking interactions ^10–13^. This elevated stability has enabled using shorter oligonucleotides to target specific sequence regions with higher mismatch discrimination ^14,15^, and has been applied in numerous contexts including gene silencing ^16,17^, molecular diagnostic probes for polymerase chain reaction ^18^, microRNA and single nucleotide polymorphism detection ^19^, increasing the specificity of CRISPR-Cas9 targeting ^20^, and viral detection ^21,22^.

In this study, we present temperature-resolved synchrotron small angle X-ray scattering (SAXS) studies quantifying the impact of LNA modifications on the thermal stability of bilayer smectic order in concentrated solutions of GDNA. Using the 48-20T-48 design (two 48 bp duplexes connected with a flexible ssDNA linker of 20 thymines, Fig. 1 and Supplementary Fig. S1) employed in our previous studies ^1,2^, we compared the melting temperature (T_m_) of duplex end-to-end stacking in the bilayers formed by GDNA constructs with either AT or A+T terminated duplex segments, where +T refers to the LNA modification of T. Our results demonstrate a new means to quantify the effect of such modifications on the strength of end-to-end stacking between DNA duplexes.

## Results and Discussion

Fig. 2 shows SAXS profiles recorded on ∼150 mM NaCl solutions of 48-20T-48 GDNA constructs with either AT - AT DNA or A+T - A+T LNA terminal base pairs and at DNA concentrations (c_DNA_) of 250, 265, and 285 mg/ml and temperatures in the range ∼7-65 °C. The GDNA constructs are otherwise identical (complete sequences are given in Supplementary Table S1). The methods relevant to sample preparation and SAXS experiments are given in Supplementary Information and an example of buffer background subtraction is shown in Supplementary Fig. S2. At lower temperatures, a sharp small angle peak is observed at wave number *q*_1_ ≈ 0.18 nm^−1^ with harmonics at *q*_2_ ≈ 0.36 nm^−1^ and (weakly) at *q*_3_ ≈ 0.54 nm^−1^. An additional, sharp wide-angle peak at *q*_*w*_ ≈ 2.2 nm^−1^ is observed in the higher concentration samples. The wave number *q*_1_ corresponds to spatial periodicity 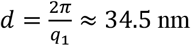 at 7 °C and c_DNA_=265 mg/ml. This is slightly greater than the ∼32 nm spacing of two 48-bp duplexes and is consistent with bilayer smectic structure considering the bilayers are separated from each other with a region where the flexible ssDNA linkers are confined. The wave number *q*_2_ corresponds to spatial periodicity 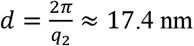 at 7 °C and c_DNA_=265 mg/ml. The peak at *q*_2_ represents the second harmonic of the bilayer phase (≈ 2*q*_1_) along with an overlapping monolayer phase. The peak at *q*_3_ represents the third harmonic of the bilayer phase. The wide-angle peak signifies positional ordering of the duplexes within the bilayers (smectic-B phase) ^3^. Assuming hexagonal packing in the smectic-B phase, *q*_*w*_ corresponds to lateral duplex-to-duplex spacing of 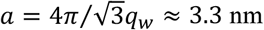 nm at 7 °C. The peak positions for *q*_1_, *q*_2_, and *q*_3_ are largely stable as a function of temperature, while *q*_*w*_ shows a modest shift to higher *q* in temperature (Supplementary Fig. S3).

**Figure 2.**
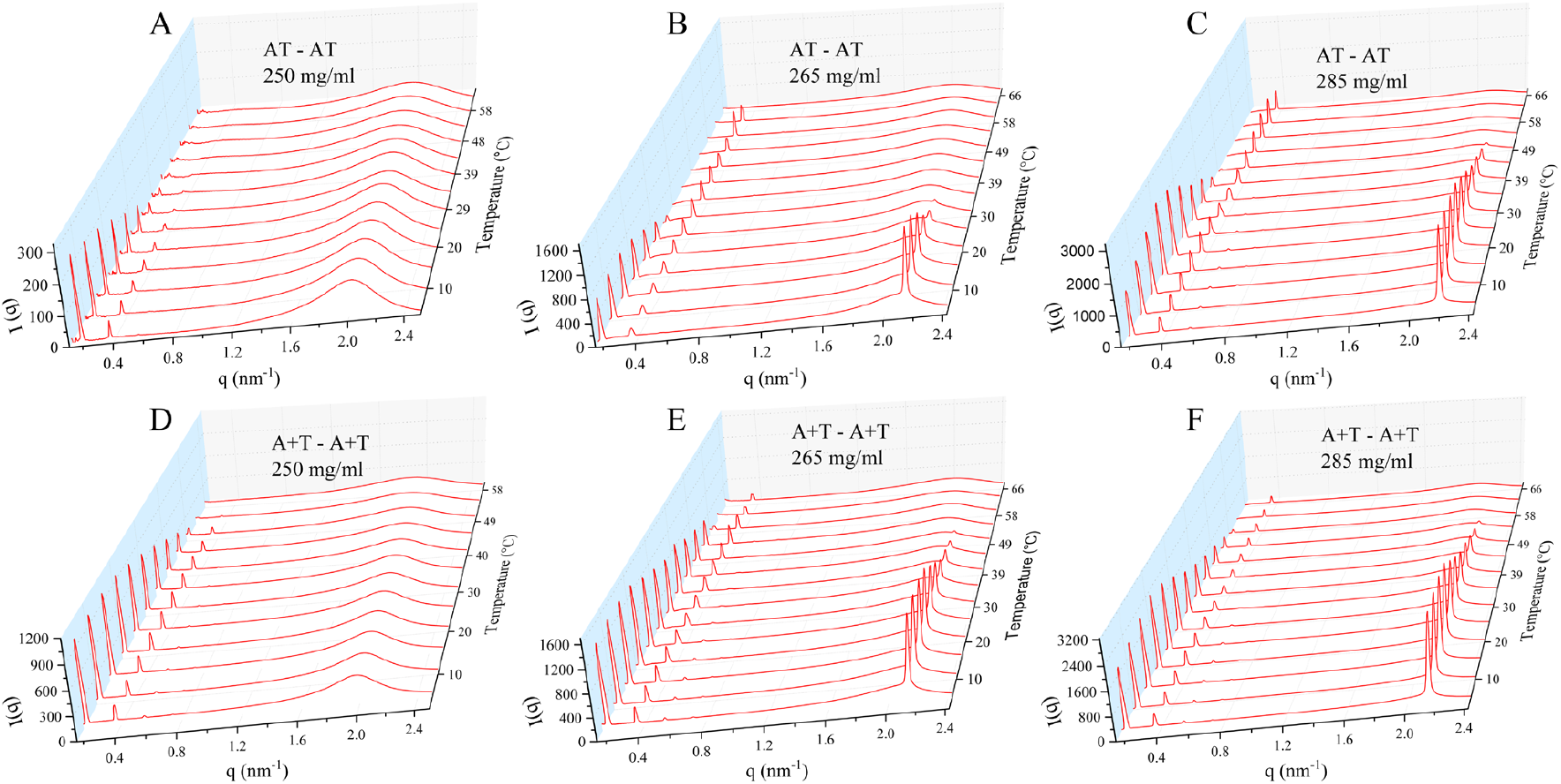
Temperature dependence of the azimuthally averaged SAXS intensity vs. scattering wavenumber *q* for GDNA samples. The data were acquired on a heating cycle with gradually increasing temperatures. (A-C) GDNA constructs with AT–AT terminal base pairs at (A) c_DNA_=250 mg/ml; (B) c_DNA_=265 mg/ml; and (C) c_DNA_=285 mg/ml. The feature at q≈0.2 nm^-1^ in the 250 mg/ml sample which persists at high temperatures is due to imperfections in subtracting the background buffer signal. (D-F) GDNA constructs with A+T - A+T terminal base pairs at (D) c_DNA_=250 mg/ml; (E) c_DNA_=265 mg/ml; and (F) c_DNA_=285 mg/ml. The bilayer order is characterized by a peak at *q*_1_ ≈ 0.18 *nm*^−1^ (and a smaller peak at *q*_3_ *≈* 0.54 *nm*^−1^) and the monolayer order with a peak at *q*_2_ *≈* 0.36 *nm*^−1^. In-plane positional order refers to a hexagonal order within these layers and is characterized by a wide-angle peak at *q*_*w*_ ≈ 2.2 *nm*^−1^.

From the data in Fig. 2, we observe that the stability of the bilayer phase increases with c_DNA_ in solutions with LNA modified duplex termination just as it does in solutions without this modification ^1,23^. However, both the bilayer order and in-plane positional order that accompanies it are stable to significantly higher temperature for constructs with LNA modification on the terminal base pairs (4). For the c_DNA_=265 mg/ml and 285 mg/ml solutions, the bilayer smectic-B phase melts at elevated temperature into a monolayer smectic-A phase (layer spacing ≈ 2π/*q*_2_, nearly half that of the bilayer phase). For the lower concentration (c_DNA_ = 250 mg/ml) solution, in which the bilayer order is not accompanied by an in-plane positional order, the smectic phase transitions into a fluid phase without sharp SAXS peaks at ∼50 °C (Fig. 2D), which is∼15 °C higher than for the AT – AT system at the same c_DNA_. This suggests that the enhanced stability of the bilayer order, enabled by the LNA modification, is by itself not adequate to generate in-plane positional order. Consistent with earlier measurements ^2^, the LNA modified constructs exhibit significant thermal hysteresis and do not completely recover the original peak amplitude upon cooling after melting of the LC phases. An example of this is shown in Supplementary Fig. S4.

Fig. 3 shows thermal melting analysis of the data in Fig. 2, following a procedure described earlier ^4^, and summarized in Supplementary Information. The melting temperatures determined from this analysis are provided in Table 1. An important aspect of this procedure involves Gaussian fitting of the peaks observed in the SAXS data and quantifying the areas under these peaks (see Supplementary Fig. S5). From this analysis, we deduce an upward shift in melting temperature (T_m_) of the bilayer structure by 14.6 °C, 17.6 °C, and 8.7 °C between constructs with or without LNA modifications at c_DNA_ = 250, 265, and 285 mg/ml, respectively. These shifts reflect the cumulative effect of the four base pairs, which would suggest each LNA termination increases the stability of the bilayer phase by 3.7 °C, 4.4 °C, and 2.2 °C per base pair.

**Table 1.** Results for the fit parameter *T*_*m*_ from the fits using a single Hill function, and for the two parameters *T*_*m*1_ and *T*_*m*2_ determined from fits using a double Hill function. The GC - GC data (bottom row only) has been adapted from reference ^4^ and provided to facilitate the discussion.

|  |  | Bilayer Order Melting |  |  | In-plane Order Melting |  |  |
| --- | --- | --- | --- | --- | --- | --- | --- |
|  |  | Single Hill Function Fit | Double Hill Function Fit |  | Single Hill Function Fit | Double Hill Function Fit |  |
|  | c<sub>DNA</sub>(mg/ml) | T<sub>m</sub>(°C) | T<sub>m1</sub>(°C) | T<sub>m2</sub>(°C) | T<sub>m</sub>(°C) | T<sub>m1</sub>(°C) | T<sub>m2</sub>(°C) |
| AT - AT | 250 | 26.7±0.5 | 20.3±1.0 | 32.8±1.6 | N/A | N/A | N/A |
| AT - AT | 265 | 30.5±0.5 | 25.1±0.2 | 34.1 | 20.9±0.4 | 15.7±0.3 | 26.6±0.4 |
| AT - AT | 285 | 39.1±0.5 | 32.7±0.3 | 45.4±0.3 | 37.2±0.8 | 29.0±0.4 | 45.2±0.3 |
| A+T – A+T | 250 | 41.3±0.9 | 32.2±1.0 | 47.8±0.9 | N/A | N/A | N/A |
| A+T – A+T | 265 | 48.1±1.5 | 36.0±0.6 | 57.5±0.3 | 33.5±0.7 | 25.0±0.6 | 43.4±0.7 |
| A+T – A+T | 285 | 48.5±1.0 | 38.1±0.6 | 56.7±0.6 | 38.7±1.0 | 28.7±0.4 | 49.7±0.4 |
| GC - GC | 280 | 64.4±0.4 | 51.7±0.2 | 67.3±0.8 | 58.6±0.2 | 47.3±0.8 | 65.6±0.6 |

**Figure 3.**
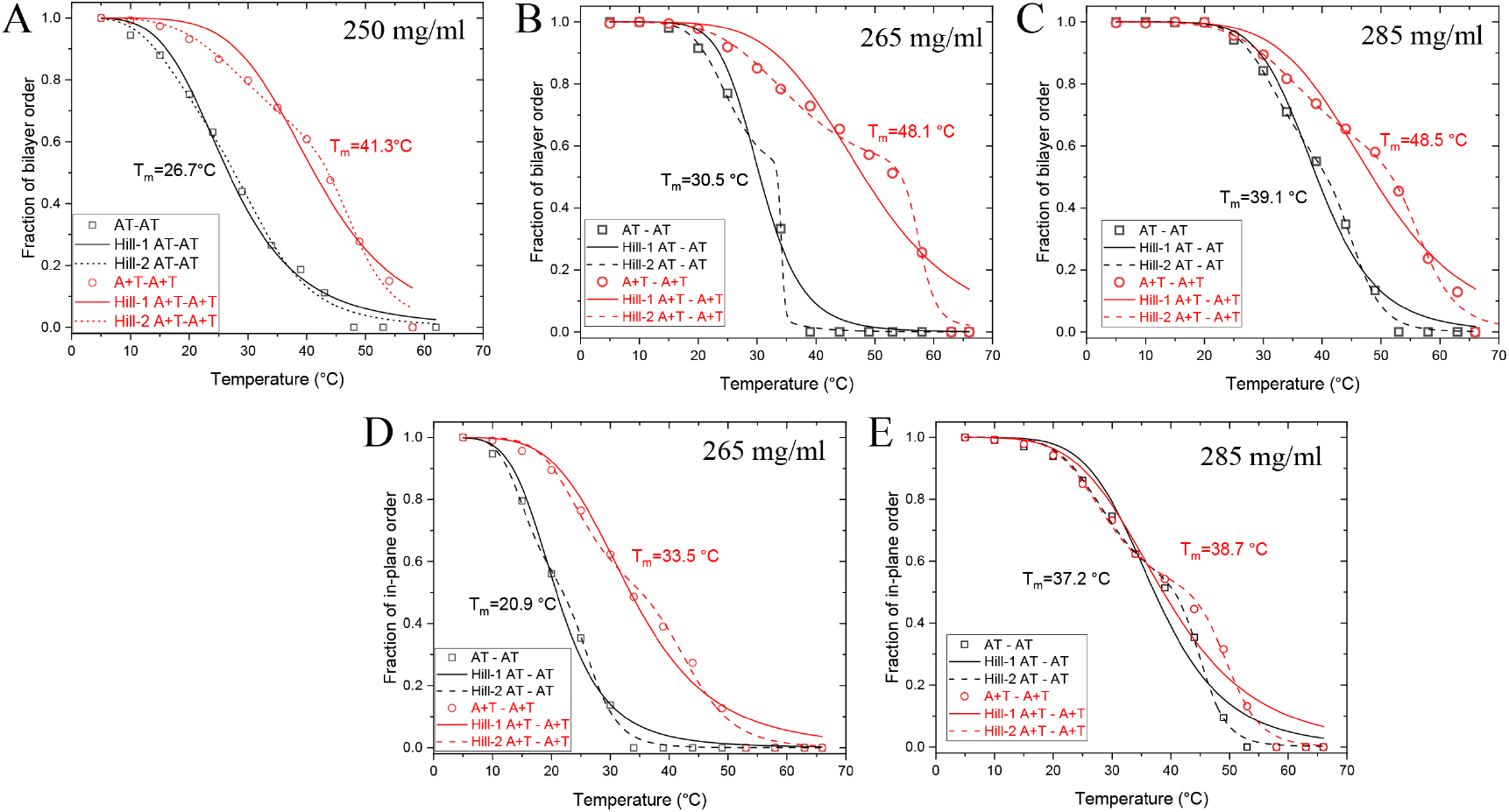
Thermal stability analysis of the SAXS data of Fig. 2. The average fractions of end-end stacked duplexes constituting the bilayer domains were determined based on the areas under Gaussian fits to the small angle peaks at *q*_*1*_ and *q*_*3*_ (specifically associated with the bilayer structure), as described in Supplementary Information. A similar analysis was performed for the sharp wide-angle peak at *q*_*w*_ (associated with in-layer ordering of the duplexes). (A-C) Fraction of bilayer order for the constructs with or without LNA modification are overlaid for 250 mg/ml in (A); 265 mg/ml in (B); and 285 mg/ml in (C). Similar analysis is performed for the in-plane positional order in (D) and (E) for c_DNA_=265 mg/ml and 285 mg/ml, respectively (in plane positional order is not observed for c_DNA_=250 mg/ml). The solid and dashed lines are fits of the data to a single and double Hill functions, respectively, from which characteristic melting temperatures *T*_*m*_ or *T*_*m*1_ and *T*_*m*2_ are determined (reported in Table 1).

The model by Fereira et al. ^10,13^ proposed the hydrogen bonding between an LNA modified AT base pair to be similar to that of a GC base pair, while the stacking interactions were similar to that of an unmodified AT base pair. Assuming this model, we would attribute the enhanced stability of the A+T – A+T bilayer smectic phase compared to that of the AT - AT construct to stronger hydrogen bonding, which reduces end-fraying and facilitates stacking of the neighboring duplexes. In a previous study, we demonstrated that adding two unpaired thymine nucleotides at the end of the duplexes completely destabilized the bilayer smectic phase at ambient temperature (25 °C) ^2^. Supplementary Fig. S6 shows base-pair opening profiles for our GDNA construct (with AT, A+T, or GC terminal base pairs) using the Fereira et al. model. The calculations show that A+T termination has a similar opening profile to that of GC termination; however, it is much more stable than an AT terminal base pair. Supplementary Fig. S7 shows a similar comparison for AT and TT, to illustrate the effect of end-fraying, which is much more likely for non-complementary bases such as TT.

Comparing the results of the current study with those of earlier work ^4^, we can estimate the contributions of stronger hydrogen bonds (which result in reduced end-fraying) and elevated stacking interactions to the enhanced stability observed for the bilayer smectic phase of the GDNA constructs with GC - GC termination relative to those with AT - AT termination. In the earlier study, we determined T_m_=64.4±0.4 °C for GC - GC terminal base pairs and T_m_=34.8±0.4 °C for AT - AT terminal base pairs in solutions of 48-20T-48 constructs with c_DNA_≈280 mg/ml. The ∼30 °C difference in stability is due to both reduced end-fraying of the four duplexes involved in formation of the bilayer phase and elevated pair-wise base stacking interactions between them. In the current study, we determined T_m_=48.5±1.0 °C for A+T – A+T terminal base pairs and T_m_=39.1±0.5 °C for AT – AT terminal base pairs at c_DNA_=285 mg/ml. We make the following two assumptions to proceed with the estimate: (i) the ∼9 °C difference in stability is due to stronger hydrogen bonds in the A+T – A+T system; and (ii) the hydrogen bonds in A+T base pair are similar in strength to those of a GC base pair (based on Ferreira *et al*. model). Given these assumptions, we estimate that the ∼30 °C difference in the stability of GC – GC and AT – AT terminations results from a cumulative effect of ∼9 °C due to stronger hydrogen bonds and ∼21 °C due to stronger stacking interactions. These would suggest base stacking of blunt duplex ends contributes approximately 2.3-fold more than hydrogen bonding of terminal base pairs to the stability of bilayer smectic phase. This is in-line with the 2-to 3-fold higher contribution of base stacking interactions to duplex DNA stability compared to hydrogen bonding (after averaging over sequence variations) ^24,25^.

## Conclusion

Our study has several important conclusions. First, we demonstrate that LC phases formed by GDNA constructs show high enough sensitivity to detect a single terminal LNA modification. The resolution of the approach is high enough to distinguish an A+T terminal base pair from AT or a GC terminal base pair based on the thermal stability of the bilayer smectic phase. Using a comparative approach, we estimated the contributions of base stacking and hydrogen bonding of the terminal bases to the stability of bilayer ordering of duplexes. Also, LNA modifications have resulted in separation of the bilayer smectic phase and the in-plane positional order over a broader temperature range (∼7-50 °C) than previously achieved. This study illustrates that LNA modification of terminal base pairs provides an additional mechanism, distinct from sequence variations of unmodified DNA, to modulate the stability of LC phases and create biomaterials with desired characteristics.

### Supporting Information Available

Sequences of DNA constructs, procedures on GDNA synthesis, SAXS measurements, thermal melting analysis, data on buffer background subtraction, temperature dependence of SAXS peak positions, data showing thermal hysteresis, data showing Gaussian fitting to SAXS peaks, computational opening profiles of DNA constructs with different terminal base pairs

## Acknowledgements

SAXS/WAXS measurements were performed on the CMS beamline (11-BM) at the National Synchrotron Light Source II, a U.S. Department of Energy (DOE) Office of Science User Facility operated for the DOE Office of Science by Brookhaven National Laboratory under Contract No. DE-SC0012704. The authors are particularly grateful to R. Li, F. Yang, and M. Fukuto for their assistance in these measurements. We thank G. Weber (Federal University of Minas Gerais, Brazil) for insightful discussions and for calculating the opening profiles for our constructs with different terminations.

