## Supporting Information for "Locked Nucleic Acid Stabilized Liquid Crystalline Phases"

Sineth G. Kodikara[a], Samuel Sprunt[a,b,]*, and Hamza Balci[a,]*
[a] Department of Physics, Kent State University, Kent, OH 44242, USA;
[b] Advanced Materials and Liquid Crystals Institute, Kent State University, Kent, OH 44242.

### Materials And Methods

**Gapped DNA (GDNA) Synthesis:** DNA oligomers (O1, O2, and O3) were purchased from Genescript (Piscataway, NJ) in polyacrylamide gel electrophoresis (PAGE) purified form. The sequences of O1, O2, and O3 for both AT – AT and the A+T – A+T constructs are given in Table S1. The A+T – A+T constructs were created by replacing the thymine (T) at the 3' end of O2 and the T at the 5'-end of O3 with LNA-modified thymine (+T). The strands containing LNA modifications are called O2+ and O3+ to distinguish them from the unmodified strands.

GDNA synthesis and sample loading into thin-walled borosilicate capillaries were carried out as described previously (1). The GDNA constructs (48-20T-48) consist of two symmetric 48-base pair duplexes connected by a 20 nt long single-stranded DNA gap of consecutive thymine bases (Fig.1A). To create the GDNA constructs, a long strand (O1, 116 nt) is annealed with two short strands (O2 and O3 for AT – AT construct or O2+ and O3+ for A+T – A+T construct, each 48 nt) which are complementary to either side of O1 except the 20T sequence in the middle. After annealing, the GDNA solutions are passed through a 50 kDa membrane filter (Amicon Ultra from Millipore) to remove incomplete constructs and diluted such that they contain ~30 mM NaCl by adding distilled and deionized water. This solution is then concentrated by centrifuging it for 12 min at 12,000 rpm (GDNA concentration increases while NaCl concentration is kept at ~30 mM). The GDNA concentration is measured with a NanodropOne Spectrometer (Thermo Fisher Scientific) and is typically in the 80-100 mg/ml range at this step. This solution is then added to quartz capillaries (2 mm inner diameter) with an open end. Water is slowly evaporated which results in increasing GDNA and NaCl concentration. The NaCl concentration is adjusted such that ~150 mM NaCl will be reached when the desired DNA concentration is achieved.

**SAXS Measurements:** SAXS measurements were carried out on beamline 11-BM at the National Synchrotron Light Source II. The incident X-ray energy was 17 keV, and the incident beam size at the sample was 0.2 × 0.2 mm. The typical acquisition time for SAXS patterns was 5 s, which did not result in visible X-ray damage to samples. A commercial hot/cold stage with Kapton film windows was used to regulate the sample temperature between 5-65 °C, which is well below the thermal melting temperature (~81 °C at the relevant DNA concentration) of the 48-bp duplex arms. We also recorded background scattering from a capillary containing pure buffer solution, which was subtracted from the data taken on the GDNA samples during processing of the SAXS patterns. A capillary filled with silver behenate powder was used to calibrate the scattering wave number ($q$) in the plane of the detector.

| Construct | Strand | Sequence (5′ to 3′) | Length(nt) |
| --- | --- | --- | --- |
| AT - AT | O1 | ACAGATGCACATATCGAGGTGGACATCACTTACGCTGAGTACTTCGAATTTTTTTTTTTTTTTTTTTTAAGCTTCATGAGTCGCATTCACTACAGGTGGAGCTATACACGTAGACA | 116 |
| | O2 | TTCGAAGTACTCAGCGTAAGTGATGTCCACCTCGATATGTGCATCTGT | 48 |
| | O3 | TGTCTACGTGTATAGCTCCACCTGTAGTGAATGCGACTCATGAAGCTT | 48 |
| A+T – A+T | O1 | ACAGATGCACATATCGAGGTGGACATCACTTACGCTGAGTACTTCGAATTTTTTTTTTTTTTTTTTTTAAGCTTCATGAGTCGCATTCACTACAGGTGGAGCTATACACGTAGACA | 116 |
| | O2+ | TTCGAAGTACTCAGCGTAAGTGATGTCCACCTCGATATGTGCATCTG+T | 48 |
| | O3+ | +TGTCTACGTGTATAGCTCCACCTGTAGTGAATGCGACTCATGAAGCTT | 48 |

**Table S1.** Sequences of the DNA oligos used to create the GDNA constructs.

**Thermal Melting Analysis:** Thermal melting analysis is performed as described in more detail previously (2). We analyzed the data in Fig. 2 by calculating the sum of the areas under the first and third order small angle peaks ($q_1$ and $q_3$) that are specifically associated with the bilayer stacking. We first fit the data in each peak to a Gaussian function of $q$ using Origin software, and then integrated the fit result over $q$ (excluding any contribution from the background scattering). Fig. S4 shows an example of such fitting. Fig. 3 shows the square root of the resulting total integrated intensity under the peaks at $q_1$ and $q_3$, normalized to its maximum value (denoted as $\sqrt{i_{1+3}}$ ) as a function of temperature for all constructs studied. The normalized quantity $\sqrt{i_{1+3}}$ represents the average fraction ($f$) of end-to-end "bound" pairs of duplexes making up the GDNA bilayers. The argument assumes that the temperature-dependence of $f$ is proportional to that of the amplitudes $\rho_n$ contributing to the density wave, $\rho = \bar{\rho} + \sum_{n \geq 1} \rho_n \cos q_n z$ ($z$ = axis normal to the layers, $\bar{\rho}$ = average density), that describes the bilayer stacking. This assumption is reasonable if attractive end-to-end interaction between duplexes is the essential mechanism driving the bilayer formation. Only the peaks at $q_1$, $q_3$, which are associated with $\rho_1$, $\rho_3$, are included in the analysis, because at elevated temperature the peaks at $q_2$, $q_4$ are partially overlapped by diffraction (at slightly higher $q$) from the developing domains of the monolayer phase.

Fig. 3 reveals that the fraction $f$ decreases within different temperature ranges that correlate with DNA concentration, and the terminal base pairs (LNA vs DNA). The dashed and solid lines are fits of the data in Fig. 3 for $\sqrt{i_{1+3}} = f$ to single and double Hill functions,[25–27] respectively. The expressions for the Hill functions are as follows:

$f(T) = 1 - \frac{T^n}{T_m^n + T^n}$ and $f(T) = 1 - \frac{1}{2}\left(\frac{T^n}{T_{m1}^n + T^n} + \frac{T^k}{T_{m2}^k + T^k}\right)$, where $f(T)$ is the average fraction of "stacked" duplexes, $T_m$, $T_{m1}$, and $T_{m2}$ are characteristic melting temperatures, and $n$ and $k$ are positive exponents. Table 1 lists these values for all the constructions we investigated. In some cases, the decrease appears to occur in two steps separated by an inflection point ("kink") in a region where the data is sparse. With limited synchrotron beam time, it was not feasible to perform measurements at higher temperature resolution, which would have avoided this kink.

### Additional Schematics

Fig. 1B shows a schematic of the bilayer smectic phase formed by a folded by GDNA. A similar bilayer phase can also be formed by the GDNA in the unfolded conformation (Supplementary Figure S1).

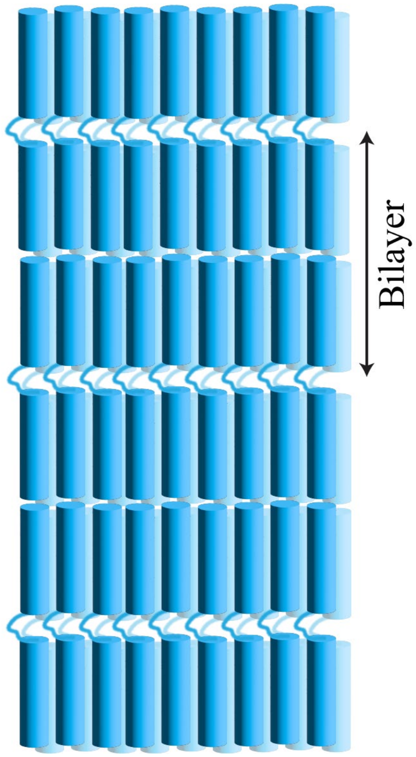


**Figure S1**. (A) Schematics of the bilayer smectic phase formed by GDNA in the unfolded conformation.

#### Buffer Background Subtraction

Figure S2 shows an example of raw sample and buffer data, in addition to final version of the data after the buffer background is subtracted from the sample.

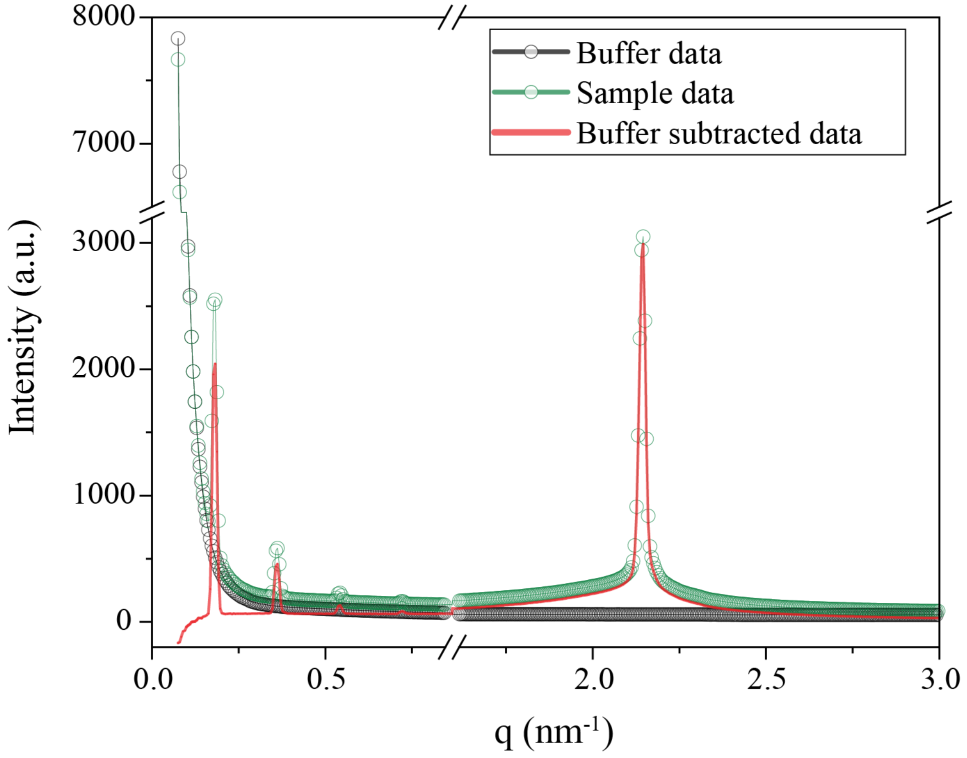


**Figure S2.** Sample data before and after the buffer background is subtracted.

### Temperature Dependence of Peak Positions

The peak positions for $q_1$, $q_2$, and $q_3$ are largely stable as a function of temperature, while $q_w$ shows a modest shift to higher $q$ in temperature (Fig. S3). This shift suggests tighter packing of the duplexes within the layers at higher temperatures.

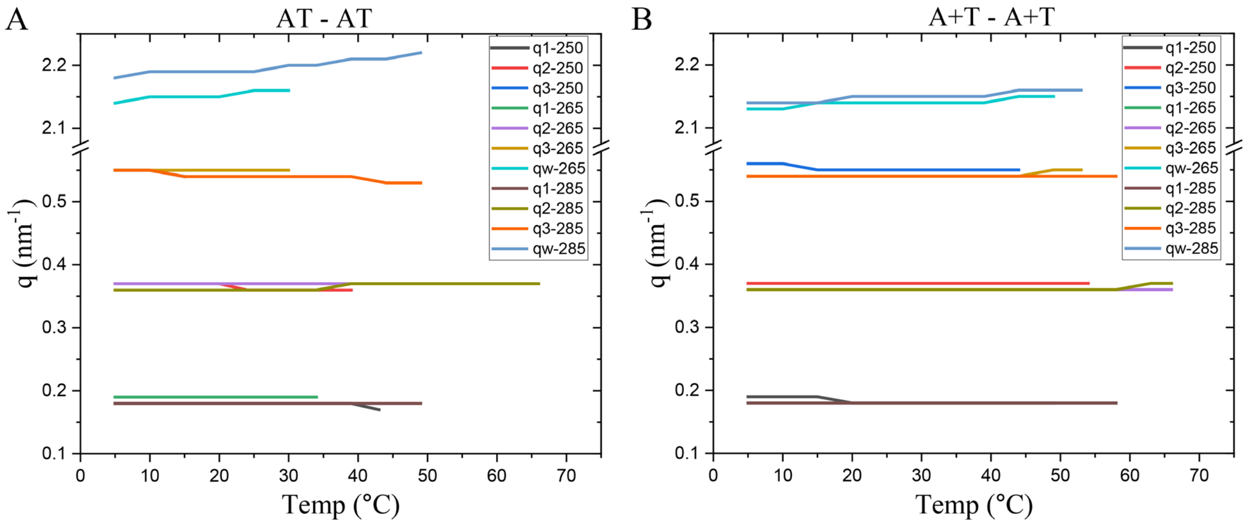


**Figure S3.** Temperature dependence of peak positions for $q_1$, $q_2$, $q_3$, and $q_w$.

### Thermal Hysteresis

The peak amplitudes in the cooling data (blue) are consistently smaller than those in heating data (red) and the level of hysteresis depends on the construct. This may be due either to the reformation time of the domains exceeding the time available for each measurement during the limited synchrotron beamtime, or to the domains reforming with a distribution of layer orientations that scatter less efficiently in the fixed incident beam and detector geometry.

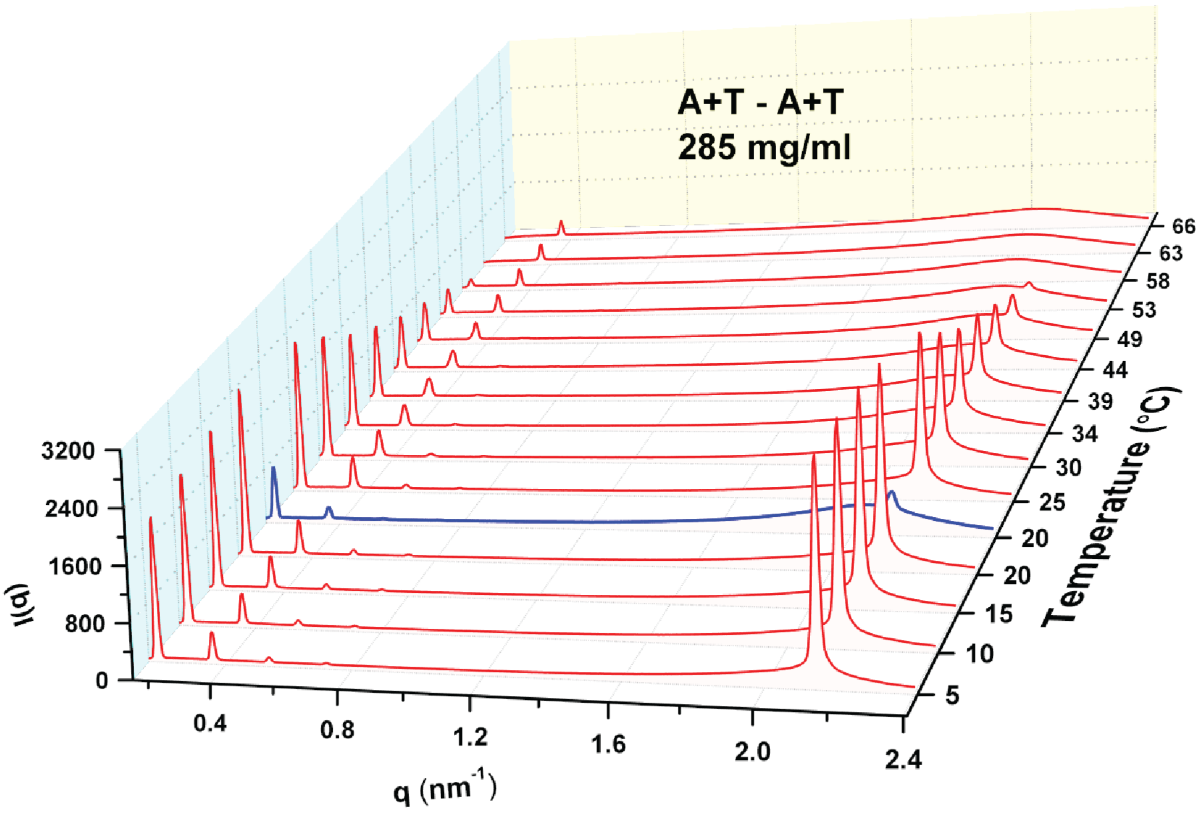


**Fig. S4.** Temperature dependence of the azimuthally averaged SAXS intensity vs. scattering wave number on heating (red traces) and subsequent cooling (blue traces) for the LNA modified GDNA construct at $c_{DNA}$=285 mg/mL.

**Gaussian Peak Fitting**

To calculate the thermal melting temperatures, the areas under each peak are calculated. First, each peak is fitted to a Gaussian function as shown in Fig. S4.

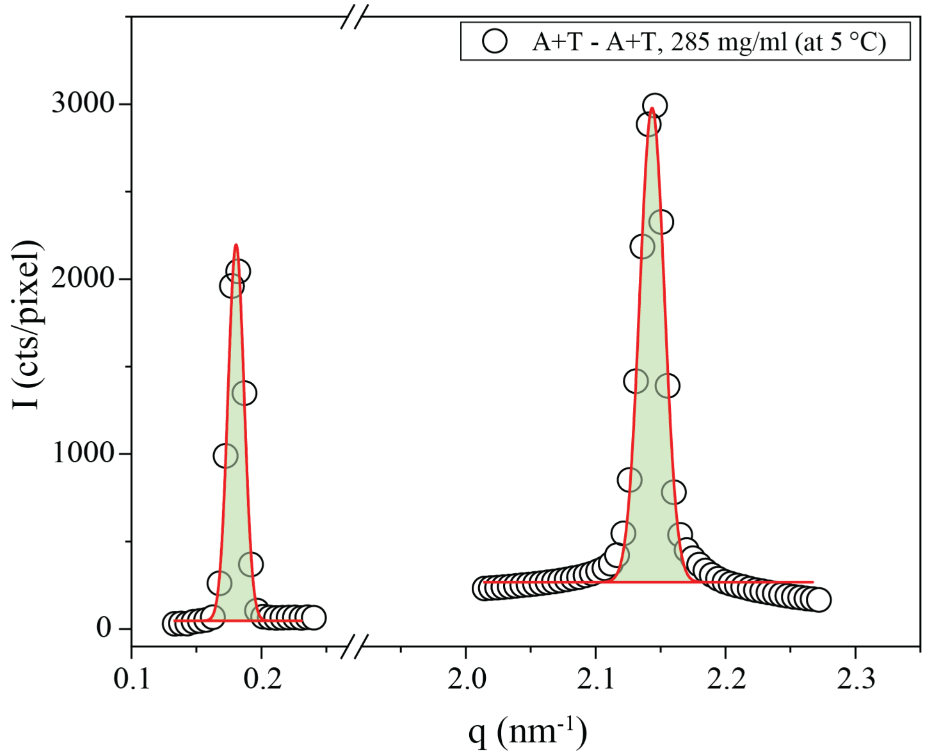


**Figure S5.** Two representative Gaussian fits are shown for the A+T - A+T, 285 mg/mL construct at 5 °C for the peaks at $q_1$ (left) and $q_w$ (right). Open symbols are data points and solid lines are the Gaussian fits. The area under each peak is shaded. The base of the Gaussian represents the background.

### Opening Profiles for GDNA with Varying Terminal Base Pairs

As discussed in the manuscript, the model by Ferreira et al. (3, 4) proposes LNA modifications to primarily enhance hydrogen bonding while stacking interactions are significantly less impacted. In this model, the hydrogen bonding between an A+T base pair is similar to that of a GC base pair, which is significantly stronger than a TA base pair. Figure S2 shows the opening profiles (a measure of base pair breaking) of the 48-20T-48 GDNA construct for different terminal base pairs. Figure S3 shows a similar comparison for TA and TT base pairs to illustrate the impact of end-fraying, which is much more likely for non-complementary bases. Both figures are based on the model by Ferreira et al.

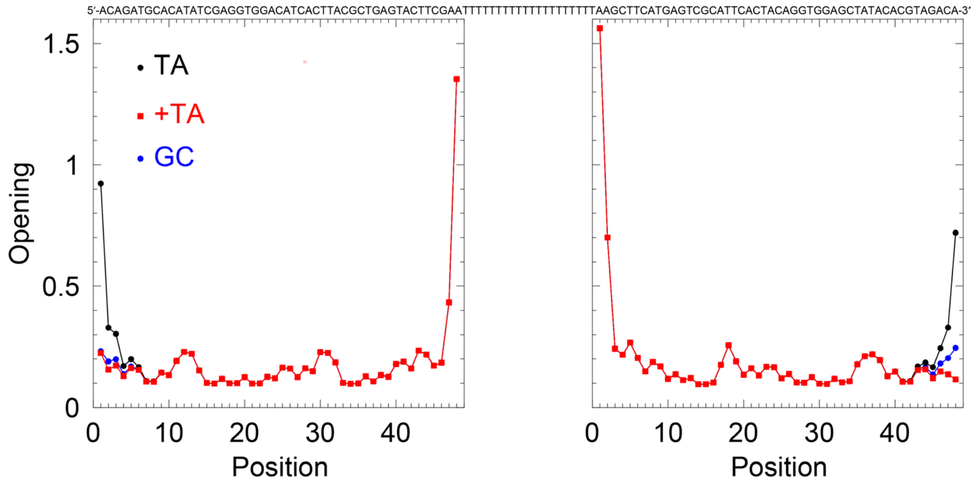


**Figure S6**. Opening profiles when terminal base pair of the GDNA construct is TA, +TA, or GC. Calculation is based on Ferreira et al. model (4).

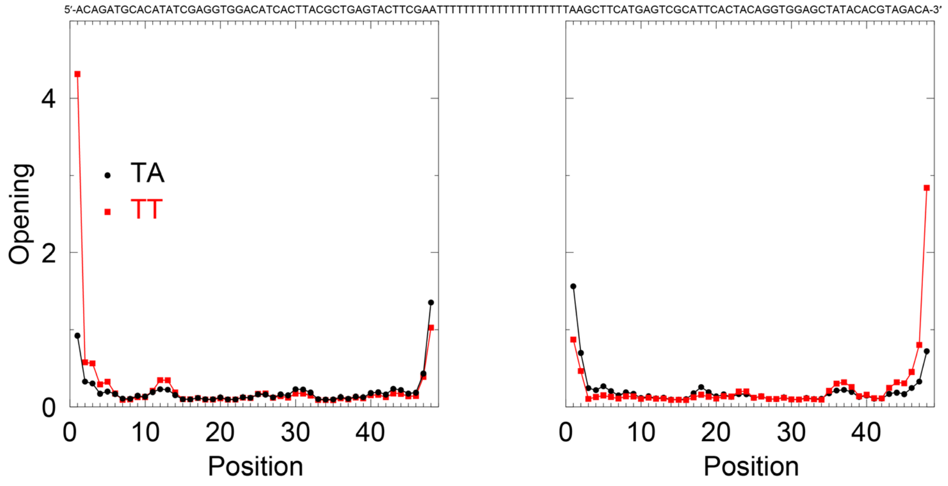


**Figure S7**. Opening profiles when terminal base pairs of the GDNA construct is TA or TT. Calculation is based on Ferreira et al. model (4).
